# Structurally Restricted Message-Passing within Shallow Architectures for Explainable Network-Level Brain Decoding on Small Cohorts

**DOI:** 10.64898/2026.03.20.713136

**Authors:** José Diogo Marques dos Santos, Maria Beatriz Ramos, Luís Paulo Reis, José Paulo Marques dos Santos, Bruno Direito

## Abstract

The application of artificial intelligence (AI) to functional magnetic resonance imaging (fMRI) has gained increasing attention due to its ability to model complex, high-dimensional brain data and capture nonlinear patterns of neural activity. However, deep learning architectures, such as Graph Neural Networks (GNNs), typically require large sample sizes to achieve stable convergence, limiting their applicability in neuroimaging contexts where data are often scarce. This challenge highlights the need for compact, data-efficient models that maintain predictive performance and interpretability.

Shallow neural networks (SNNs) have demonstrated robustness in low-sample settings but commonly rely on region-level features that treat brain areas independently, overlooking the brain’s intrinsically network-based organization. To address this limitation, we propose a structurally constrained message-passing framework that integrates diffusion tensor imaging (DTI)-derived structural connectivity with region-level fMRI signals within a shallow architecture. This approach enables network-level modeling while preserving the stability and data efficiency of SNNs.

The method is evaluated on 30 subjects performing a Theory of Mind (ToM) task from the Human Connectome Project Young Adult dataset. A baseline SNN achieved global accuracies of 88.2% (fully connected), 80.0% (pruned), and 84.7% (retrained), while the proposed model achieved 87.1%, 77.6%, and 84.7%, respectively. Although structural constraints led to a more pronounced performance decrease after pruning, retraining restored accuracy to baseline levels, demonstrating that biological constraints can be incorporated without compromising predictive validity.

Model interpretability was assessed using SHAP (Shapley Additive Explanations). While the baseline model primarily identified isolated regions as key contributors, the proposed framework revealed distributed, structurally coherent networks as the main drivers of classification. These networks showed correspondence with established ToM regions, including the temporo-parietal junction, superior temporal sulcus, and inferior frontal gyrus. Importantly, the findings suggest that groups of moderately informative regions can collectively form highly relevant subnetworks.

Overall, the proposed framework achieves competitive performance in a limited dataset while incorporating graph-inspired message passing into a shallow architecture. Its explainability provides insight into how structurally constrained networks support stimulus-driven responses in ToM and demonstrates potential for investigating network dysfunction in disorders such as Alzheimer’s disease, ADHD, autism spectrum disorder, bipolar disorder, mild cognitive impairment, and schizophrenia.

## 1 Introduction

Understanding how the brain produces sophisticated cognitive and social behavior remains a major issue in neuroscience. Functional magnetic resonance imaging (fMRI) has revolutionized the mapping of neural activity. However, most machine learning approaches to fMRI data fail to account for the anatomical pathways that connect brain regions and treat their data as isolated entities (Mohammadi & Karwowski, 2025). This is a serious shortcoming, as there is converging evidence that considers a range of neurological and psychiatric diseases, from Alzheimer’s disease to schizophrenia, as products of large network connectivity-related syndromes instead of isolated regional dysfunction (Bassett & Sporns, 2017; Du et al., 2018; Repple et al., 2023). In parallel, the opacity of artificial neural network classifiers remains a problem in their translation into neuroscientific knowledge (Farahani et al., 2022). Additionally, introducing graph neural networks (GNNs) to model the brain’s non-Euclidean nature requires large amounts of data for proper training, which is only available in extensive and costly studies. Addressing these gaps requires developing analytical frameworks that are simultaneously constrained by brain anatomy yet capable of providing principled explanations in a frugal data frame. The present study introduces a framework that combines diffusion-based structural connectivity into a shallow neural network classifier using message-passing, while analyzing its classifications using Explainable Artificial Intelligence approaches within the context of a Theory of Mind approach.

Artificial neural networks (ANNs) can extract meaningful patterns from complex datasets, producing models with high predictive accuracy when properly trained and good generalization when the input data are sufficiently representative (Haykin, 2009). In parallel, Functional Magnetic Resonance Imaging (fMRI) has become one of the most widely used techniques for studying the functioning human brain, producing four-dimensional maps of neural activity across space and time. Applications of fMRI range from investigating mental disorders to brain-decoding approaches aimed at understanding cognitive processes. Early reviews of machine learning methods for fMRI classification highlighted important methodological concerns regarding the use of ANNs in this context (Pereira et al., 2009). In particular, two key issues were emphasized: (1) whether the improvement in classification performance justifies the increased complexity of ANN models, and (2) the “black box” nature of such models, which complicates their explanation and interpretation.

Furthermore, in an opinion article, Wen et al. (Wen et al., 2018) discuss the use of diverse ANN architectures for diagnosing cognitive impairments from fMRI data. Although they highlight the potential of deep models to advance mental health research, they emphasize a critical limitation: typical fMRI datasets are relatively small, increasing the risk of overfitting in complex Deep Neural Networks (DNNs). These constraints necessitate architectures that balance predictive performance and robustness in low-data regimes. Nevertheless, ANNs remain highly relevant in Social Neuroscience, where they are used to (i) predict behavior from brain activity, (ii) quantify naturalistic social stimuli and interactions, and (iii) generate cognitive models of social brain function (Sievers & Thornton, 2024).

More recent work has begun to address these concerns by developing procedures specifically tailored to the characteristics of fMRI data. Such approaches have demonstrated that ANN-based models can produce results consistent with established findings in Neuroscience, including in cognitive paradigms related to Theory of Mind (ToM) (Marques dos Santos & Marques dos Santos, 2022, 2023; Marques dos Santos et al., 2025). However, these procedures have primarily focused on region-level analyses in which each brain region is treated independently. As a result, they identify which regions contribute to classification but do not address how these regions interact as components of anatomically organized brain networks.

This limitation is particularly relevant in the context of neurological and psychiatric disorders. Increasing evidence suggests that many such conditions are better understood as disorders of large-scale network dysfunction rather than isolated regional abnormalities. For example, Alzheimer’s disease (AD), Mild Cognitive Impairment (MCI), Attention Deficit Hyperactivity Disorder (ADHD), Autism Spectrum Disorder (ASD), Bipolar Disorder, and Schizophrenia have increasingly been conceptualized as dysconnectivity syndromes involving disruptions of large-scale brain networks (Du et al., 2018). Consequently, understanding not only which brain regions are engaged but also how they interact within biologically plausible networks is essential for advancing translational applications. More broadly, advances in artificial intelligence applied to medical imaging have been highlighted as potentially contributing to global health initiatives such as United Nations Sustainable Development Goal 3, which aims to ensure good health and well-being for people of all ages (Chauhan et al., 2024). By improving the analysis and interpretation of neuroimaging data, ANN-based methods may, therefore, contribute to both fundamental neuroscience research and clinical innovation. Importantly, the usefulness of such models is likely to increase as their design more closely reflects the brain’s biological organization.

To address the limitations of region-level analysis, the present study integrates diffusion-based structural connectivity derived from Diffusion Tensor Imaging (DTI) into a shallow neural network (SNN) classifier through a message-passing mechanism. Message-passing, commonly used in graph-based architectures, enables information to propagate along anatomically defined pathways. In the proposed framework, this mechanism allows signals from individual brain regions to be combined according to structural connectivity patterns. Importantly, it is implemented within a shallow architecture designed to perform reliably under the relatively small sample sizes typical of task-based fMRI studies. This design enables classification at the level of structurally defined networks while preserving the stability advantages associated with SNNs.

To address the second concern raised above, the “black box” nature of ANN models, the proposed framework is combined with interpretability and explainability methods. Even in relatively compact models such as the present SNN, which contains 3,620 connections, the network’s internal operations are too complex to be readily interpreted by direct inspection. Explainable Artificial Intelligence (XAI) aims to overcome this challenge by providing tools that clarify how machine learning models reach their decisions. According to Doran et al. (Doran et al., 2018), four key challenges characterize explainability in dense ANN models: opacity, where the internal mechanisms are inaccessible to users; interpretability, where the mechanisms can be analyzed mathematically; comprehensibility, where symbolic outputs enable human-understandable explanations; and full explainability, where automated reasoning generates self-contained explanations. XAI methods may be applied to increase the transparency of ANN models, enabling knowledge extraction from fMRI data in the context of a typical cognitive task (Roscher et al., 2020).

The cognitive paradigm investigated in this study originates from the field of Social Neuroscience, which examines the neural bases of social cognition. Over the past two decades, advances in fMRI have greatly accelerated research in this domain (Adolphs, 2001; Decety & Keenan, 2006; Norman et al., 2010; Saxe, 2006). Among the most extensively studied constructs in social neuroscience is the Theory of Mind, also referred to as cognitive empathy or mentalizing (Shamay-Tsoory et al., 2009). This concept describes the human ability to attribute beliefs, intentions, and goals to others during social interactions (Singer & Tusche, 2014). Such attribution of mental states plays a central role in successful social behavior. Although the extent to which ToM abilities are uniquely human remains debated (Call & Tomasello, 2008; Premack & Woodruff, 1978; van Mourik et al., 2024; Vilone & Longo, 2021), the concept remains fundamental for understanding the neural mechanisms underlying social cognition. The data used in the present study were acquired using a Theory of Mind paradigm that replicates the classic experiment introduced by Fritz Heider and Marianne Simmel in 1944 (Heider & Simmel, 1944), in which participants inferred social interactions from the movement of simple geometric shapes.

In sum, the present study aims to further develop on previous region-level approaches by investigating whether incorporating structural connectivity into an ANN framework can enable biologically meaningful network-level brain decoding while preserving classification performance. Specifically, diffusion-based connectivity is integrated into a shallow neural network via a message-passing mechanism, enabling signals from individual regions to be combined along anatomically plausible pathways. This network-level representation is then examined using methods from Explainable Artificial Intelligence to determine which structurally defined subnetworks contribute to classification in a Theory-of-Mind paradigm. In doing so, the study addresses several key questions: whether structurally constrained models can achieve predictive performance comparable to region-based approaches, whether such models yield explanations consistent with established findings in Social Neuroscience, and whether integrating anatomical connectivity can improve the biological interpretability of ANN-based analyses of Functional Magnetic Resonance Imaging data. Together, these objectives aim to evaluate whether combining structural constraints with interpretable machine learning provides a viable framework for studying both cognitive processes and network-level brain organization.

## 2 Methods

### 2.1 Dataset

The dataset comprised the social paradigm of 30 subjects from the Human Connectome Project Young Adult Dataset (HCP Young Adult; https://www.humanconnectome.org/study/hcp-young-adult; accessed on 16 September 2024), specifically subjects 100307 to 124422 from the 100 Unrelated Subjects subset (Elam et al., 2021; Van Essen & Glasser, 2016; Van Essen et al., 2013). The task consists of 20-second videos showing moving geometric shapes, whose motion is either random (rnd) or depicts interactions that require mentalizing (mnt) (Heider & Simmel, 1944). Participants classify each video as random (rnd), mentalizing (mnt), or unsure, or may provide no response. The videos are presented in their entirety regardless of when a response is given. Only correctly classified trials are included in the analysis. Each participant completes two runs with different scanner phase-encoding directions: left-to-right (LR) and right-to-left (RL). Each run includes five stimuli (two from one condition and three from the other, counterbalanced across runs). Consequently, each participant evaluates five stimuli per condition, resulting in a total of ten stimuli.

### 2.2 Feature Extraction

For SNN feature extraction, the brain is parcellated using the Human Connectome Project Multimodal Parcellation Atlas (HCP-MMP1.0) (Glasser et al., 2016). Each subject’s BOLD (blood oxygen level-dependent) image is masked with the 360 regions defined in the atlas, and the voxels’ time series within each region are averaged, yielding 360 regional mean time series per session. The complete list of all 360 regions, including their IDs, names, and locations, is provided in Supplementary Table 1. The time series are segmented according to the stimulus sequence, and the 7^th^ to 9th time points after stimulus onset are averaged to define a single feature per region for each instance. The time points are chosen based on a canonical double-gamma Hemodynamic Response Function (HRF) that models the signal lag relative to brain activation. As the TR is 0.72 seconds, the 7th through 9th time points correspond between 5.04 and 6.48 seconds after stimulus onset. Input matrices are then constructed and standardized across all features, with standardization performed separately for the training and testing sets to avoid data leakage.

### 2.3 Model Architecture

#### 2.3.1 Structural Connectivity Matrix

Structural connectivity is derived from a previously published HCP-MMP1.0–based group connectome (Baker et al., 2018; Briggs, Conner, Baker, et al., 2018; Briggs, Conner, Rahimi, et al., 2018; Briggs, Conner, Sali, et al., 2018a, 2018b; Briggs, Rahimi, et al., 2018; Conner, Briggs, et al., 2018a, 2018b; Conner, Briggs, Sali, et al., 2018; Sali et al., 2018), obtained through anatomy-driven deterministic tractography on diffusion-weighted images from 10 subjects using Generalized Q-Sampling Imaging (ratio = 1.25; angular threshold = 45°), with manual seeding of established anatomical pathways. Connectivity is defined by the presence of streamlines between regions, resulting in a 360 × 360 binary matrix containing 2,100 connections. A diagonal term is added to the matrix to preserve intra-regional contributions during message-passing. The complete structural connectivity matrix is available in Supplementary Table 2.

#### 2.3.2 Message-Passing Mechanism

The message-passing mechanism constitutes the functional basis of Graph Neural Networks (GNNs). It iteratively updates the representation of each node as a function of the representations of its neighboring (connected) nodes. However, GNNs are typically deep architectures that require large amounts of data to ensure convergence to a solution with acceptable performance. In the present study, only a single message-passing step is performed. Mathematically, this operation corresponds to multiplying the signal matrices (training and testing) by the structural connectivity matrix. Thus, the signal of each network corresponds to the sum of the signals from all structurally connected regions. This procedure is defined as:

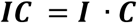

where *I* denotes the signal matrix obtained during feature extraction (an *n* × 360 matrix), *C* represents the structural connectivity matrix (a 360 × 360 matrix), and *IC* is the resulting matrix after one message-passing step according to connectivity matrix *C* (*n* × 360). A degree-based correction factor is subsequently applied by dividing the aggregated signal by the number of connections (i.e., the sum of all non-zero entries in the corresponding column of matrix *C*), yielding the average network signal. This correction mitigates signal dilution, which may occur when only a subset of regions within a network is recruited. In such cases, summing activation-related signal peaks with baseline activity can reduce the contrast between the activated and non-activated states of the network.

#### 2.3.3 SNN Classifier

To ensure consistency with previous studies and, therefore, discuss comparable results, the fully connected SNN comprises 360 input nodes (one per network), 10 hidden nodes arranged in a single hidden layer, and 2 output nodes (one for each class in the paradigm design). The network architecture is implemented using the AMORE library (version 0.2-15) (Limas et al., 2014), R (version 4.5.1), and RStudio (version 2025.05.1.513). The activation function applied to the hidden layer is the hyperbolic tangent (“tansig”), which produces outputs in the range]−1, 1[ and is centered at zero. The output layer uses the sigmoid activation function, which outputs values in the range]0, 1[. **Figure 1** illustrates a simplified schematic representation of the classifier architecture. The message-passing step occurs between the first two layers in the schema, with the second layer (the output of the message-passing mechanism) serving as the input to the SNN classifier.

**Figure 1.**
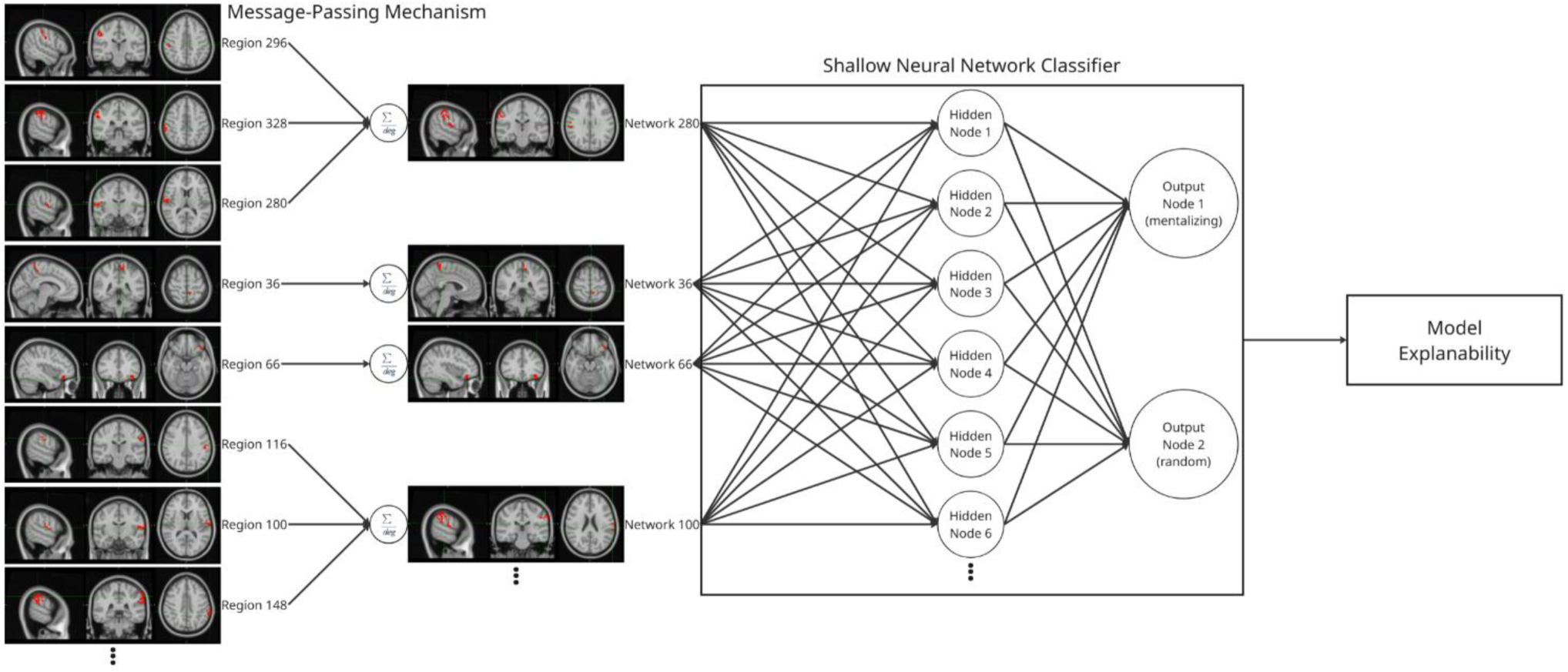
Message-passing mechanism on the SNN classifier. The connection between the first two layers (from regions to networks) is the message-passing mechanism. The following layers are the Shallow Neural Network classifier, and lastly, the model explainability steps are on the right.

### 2.4 Training, Pruning, and Retraining

Participants are randomly divided into training and testing sets. The training set includes 20 subjects (168 instances: 98 mnt and 70 rnd), while the testing set includes 10 subjects (85 instances: 45 mnt and 40 rnd). Training is performed using a grid search over the learning rate and momentum parameters. Each parameter pair is evaluated 100 times to account for variability due to random weight initialization. The best-performing pair is then used to train 50,000 networks that differ only in their initial weights, resulting in the fully connected network. The first stage of the pruning procedure consists of ranking all possible path-weights. The concept of “path-weights” (Marques dos Santos & Marques dos Santos, 2022, 2023) refers to the product of the weights along a path from a given input node to a given output node through a single hidden node, as the present architecture includes only one hidden layer. Formally, a path-weight is defined as:

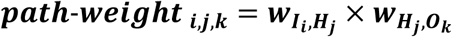

where ***w***_***Ii***,***Hj***_ is the weight between the input node, ***I***_***i***_, and the hidden node, ***H***_***j***_; and ***w***_***Hj***,***ok***_ is the weight between the hidden node, ***H***_***j***_, and the output node, ***o***_***k***_. Therefore, ***path-weight_i,j,k_*** is the product of the weights found in the path from the input ***I***_***i***_ to output ***o***_***k***_, passing by the hidden node, ***H***_***j***_. A path-weight with a magnitude close to zero tends to attenuate the contribution of the input signal along that path and may therefore be pruned due to its low relevance. Conversely, a path-weight with a larger magnitude, whether positive or negative, exerts a stronger influence on classification and is therefore retained. The distribution of path-weights typically exhibits two elbow points, indicating that only a limited number of paths effectively transmit information from the input layer to influence the model output. The elbow points are computed using the “pathviewer” (R package) (Baliga et al., 2023). Path-weights lying between the positive and negative elbow points are pruned. A more detailed description of this procedure is provided elsewhere (Marques dos Santos & Marques dos Santos, 2024). **Figure 2** shows the positive and negative arms of this distribution of the ranked path-weights.

**Figure 2.**
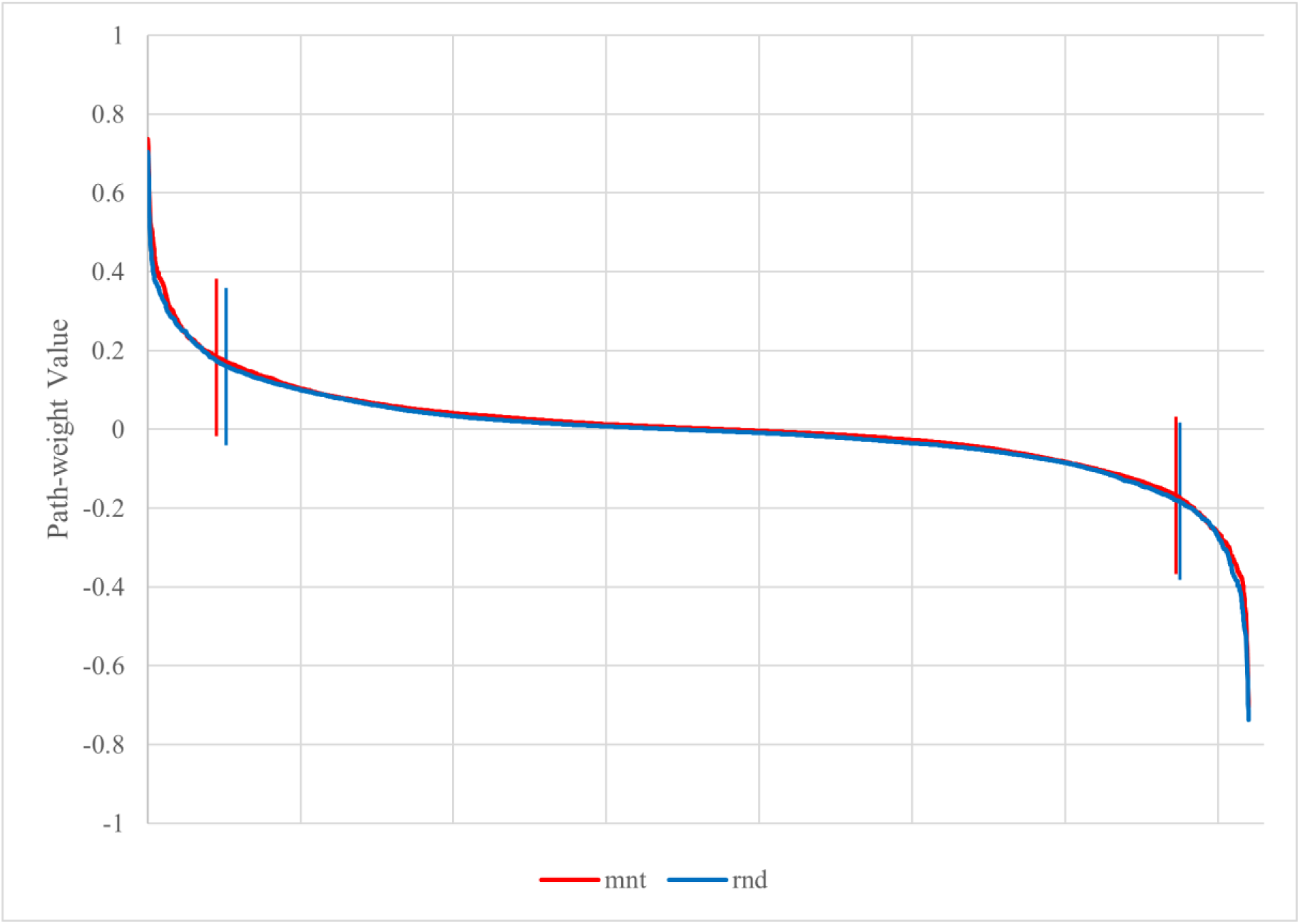
Path-weights plotted per output in decreasing order for each stimulus category. Both plots have a tilde shape, meaning that most paths have little influence on the computation (because they are close to zero and tend to cancel the signal in the multiplication). Just a few have weights that move away from nullity. The latter are retained in the pruning process. Vertical lines signal the elbow points, which correspond to: mnt (red) 226 and 3362, rnd (blue) 257 and 3375.

The resulting pruned network is then retrained to update the weights according to the new architecture. At this stage, a grid search is performed again to identify the best hyperparameters (learning rate and momentum). However, only one network is trained for each parameter pair, since the initial weights are deterministic and no longer randomly initialized. This step produces the retrained network.

### 2.5 Explaining

Shapley values are defined by:

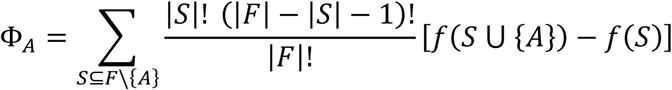

where *S* is the set of inputs to the SNN, with the exception of input *A*, which is the target of evaluation about its importance/contribution to the classification. The term [*f*(*S* ⋃ {*A*}) − *f*(*S*)] is essential for comprehending the significance of the Shapley values, as it compares two situations: the full set of inputs, i.e., *f*(*S* ⋃ {*A*}), and the situation where the targeted input is removed from the calculations, i.e., *f*(*S*). Two groups of results may occur:

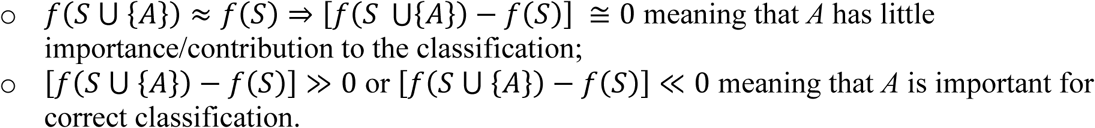

The computation of Shapley values is performed using the retrained network in Python with the SHAP library (https://github.com/shap/shap; accessed on 06 February 2026). SHAP (SHapley Additive exPlanations) provides a framework for interpreting model predictions by attributing contributions to individual features. Because the original network is implemented in R using the AMORE package, the model must first be translated to Python using PyTorch (https://pytorch.org/; accessed on 06 February 2026). To reproduce the original architecture, a custom activation function is implemented in Python to mimic the tansig activation used in AMORE. A network with the same architecture as the original model is first initialized in PyTorch, consisting of 360 input nodes, one hidden layer with 10 hidden nodes using the custom tansig activation function, and two output nodes with the sigmoid activation function. The network is initially created with randomly generated weights. The weights from the retrained network are then exported from R and imported into the PyTorch model in the appropriate format, replacing the randomly initialized weights. This procedure yields an exact replication of the network originally trained in R. The explanation stage is subsequently performed using the SHAP library. An explainer is created using the DeepExplainer function, which is based on an enhanced version of the DeepLIFT algorithm (Shrikumar et al., 2017). As shown in (Lundberg & Lee, 2017), this method approximates the conditional expectations of Shapley values by integrating over a large number of samples, such that the sum of the SHAP values equals the difference between the expected model output and the actual model prediction. The SHAP analysis uses these approximations of Shapley values to rank the features according to the change they produce in the model’s decision, thereby quantifying their importance in the SNN’s classification process. This procedure yields one value for each feature, for each instance, and for each class. These values can then be averaged in absolute magnitude to obtain a global ranking of feature importance. Furthermore, examining the relationship between the SHAP value and the corresponding input value reveals whether a feature has a positive or negative influence on the model’s decision. Specifically, this analysis indicates whether higher input values drive the prediction toward a given class or contribute to rejecting classification into that class.

In this study, the SHAP analysis identifies the most important inputs (brain networks), i.e., ranks highly the brain networks whose putative absence would mostly impact a drop in accuracy. However, these must be compared to established neuroscientific knowledge. To establish the baseline for this comparison a GLM-based statistical parametric map of a similar mentalizing condition using Frith-Happé animations is extracted from the literature (Dureux et al., 2023). To obtain a quantitative measure of the overlap between this established baseline and the identified relevant networks from the SHAP analysis, the overlap coefficient is used. For this study, the overlap is mathematically given by:

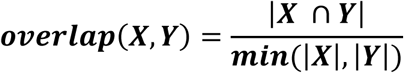

where ***overlap (X,Y)*** is the overlap coefficient between maps ***X*** and ***Y***, |***X***| and |***Y***|are the total number of voxels in each map, and |***X*** ∩ ***Y***| is the number of voxels common to both maps. In practice, it provides a quantitative measure of how much of the identified networks also appear in the baseline GLM-based map.

### 2.6 Cohen’s D

Cohen’s D is an effect size measure based on differences between means. Effectively, it measures the distance between two sets of data points. Thus, Cohen’s D value closer to 0 means both sets of data points are close together, while a higher or lower (more negative) value means they are far apart.

Mathematically, Cohen’s D is given by:

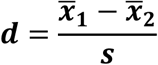

where ***d*** is the Cohen’s D value, ***x̅***_***1***_ is the mean of the first set of data points, ***x̅***_***2***_ is the mean of the second set of data points, and ***s*** is the pooled standard deviation.

Effectively, in this study, this value is calculated separately for mentalizing and random instances and compared across regions within each network. Networks identified during the SHAP analysis are important for the model to differentiate between classes. However, it provides no information regarding the contribution of each region within each network.

## 3 Results

### 3.1 Region Level Classification (baseline)

First, the regional-level classification corresponds to the model reported in the literature (Marques dos Santos et al., 2025). It is trained on the same dataset and uses identical preprocessing and feature extraction procedures. Importantly, in the original study, the explanations produced by this analysis were examined and shown to be consistent with established findings in Neuroscience. Therefore, it provides a robust baseline against which the proposed message-passing mechanism can be compared. The performance of the three network configurations is summarized in **Table 1**, which reports confusion matrices, local precisions as well as both local and global accuracies for the fully connected, pruned, and retrained networks. The initial global accuracy is 88.2%, which decreases to 80.0% after pruning. Following retraining, the global accuracy increases to 84.7%. A detailed analysis of this model is provided in the original study.

**Table 1.** Confusion matrices, precisions, and accuracies of the fully connected, pruned, and retrained networks obtained for the region-level classifier in (Marques dos Santos et al., 2025).

| Network | Stimulus |  | Prediction |  | Total |
| --- | --- | --- | --- | --- | --- |
|  |  |  | mnt | rnd |  |
| Fully Connected Network | Input | mnt | 43 | 2 | 45 |
|  |  | rnd | 8 | 32 | 40 |
|  | Total |  | 51 | 34 | 85 |
|  | Accuracy (%) |  | 95.6 | 80 | 88.2 |
|  | Precision (%) |  | 84.3 | 94.1 |  |
| Pruned Network | Input | mnt | 41 | 4 | 45 |
|  |  | rnd | 13 | 27 | 40 |
|  | Total |  | 54 | 31 | 85 |
|  | Accuracy (%) |  | 91.1 | 67.5 | 80 |
|  | Precision (%) |  | 75.9 | 87.1 |  |
| Retrained Network | Input | mnt | 41 | 4 | 45 |
|  |  | rnd | 9 | 31 | 40 |
|  | Total |  | 50 | 35 | 85 |
|  | Accuracy (%) |  | 91.1 | 77.5 | 84.7 |
|  | Precision (%) |  | 82 | 77.5 |  |

### 3.2 Network-Level Classification

Overall, the proposed network-level classification approach yields highly comparable results. These findings are summarized in **Table 2**, which reports the confusion matrices, local precisions, and both local and global accuracies for the fully connected, pruned, and retrained networks. The numbers of correct predictions are 74, 66, and 72, corresponding to global accuracies of 87.1%, 77.6%, and 84.7%, respectively. As observed previously, pruning leads to a reduction in performance that is largely recovered after retraining. In the message-passing model, the initial accuracy (fully connected network) is slightly lower than in the baseline model. The performance drop following pruning is more pronounced, yet the final accuracy after retraining matches the baseline performance.

**Table 2.** Confusion matrices, precisions, and accuracies of the fully connected, pruned, and retrained networks obtained for the network-level classifier.

| Network | Stimulus |  | Prediction |  | Total |
| --- | --- | --- | --- | --- | --- |
|  |  |  | mnt | rnd |  |
| Fully Connected Network | Input | mnt | 43 | 2 | 45 |
|  |  | rnd | 9 | 31 | 40 |
|  | Total |  | 52 | 33 | 85 |
|  | Accuracy (%) |  | 95.6 | 77.5 | 87.1 |
|  | Precision (%) |  | 82.7 | 93.9 |  |
| Pruned Network | Input | mnt | 38 | 7 | 45 |
|  |  | rnd | 12 | 28 | 40 |
|  | Total |  | 50 | 35 | 85 |
|  | Accuracy (%) |  | 84.4 | 70 | 77.6 |
|  | Precision (%) |  | 76 | 80 |  |
| Retrained Network | Input | mnt | 43 | 2 | 45 |
|  |  | rnd | 11 | 29 | 40 |
|  | Total |  | 54 | 31 | 85 |
|  | Accuracy (%) |  | 95.6 | 72.5 | 84.7 |
|  | Precision (%) |  | 79.6 | 93.5 |  |

Regarding local accuracies, performance for the mentalizing (mnt) class remains consistently higher than for the random (rnd) class across all stages of network development. However, unlike the baseline model, where pruning predominantly affects the rnd condition, the message-passing model exhibits a more balanced impact of pruning across both classes. The reduction in local accuracy is slightly greater for mnt (−11.2%) than for rnd (−7.5%). In the retrained network, the accuracy for the rnd class decreases relative to the baseline model (77.5% in the baseline vs. 72.5% in the message-passing model), whereas accuracy for the mnt class increases (91.1% in the baseline vs. 95.6% in the message-passing model). This pattern suggests that the message-passing model is particularly sensitive to mentalizing-related network patterns, potentially providing meaningful insights into stimulus-induced network recruitment associated with Theory of Mind. Local precision initially follows the same pattern observed in the baseline model: it is higher for the rnd class than for the mnt class, decreases after pruning, and increases again following retraining. However, in contrast to the baseline model, the precision for the rnd class remains higher than for the mnt class, even in the retrained network.

### 3.3 SHAP Values

To maintain a manageable interpretation of the results, only the top 20 highest-ranked SHAP values per category are considered. Because the task is a binary classification problem, the ordering of the input features by the mean absolute value of the estimated Shapley values is identical across both categories. **Figure 3** displays the beeswarm plots for the top 20 input features in each category. The complete beeswarm plots for all 360 networks, including ones that are pruned during model development, for each category, are reported in Supplementary Figure 1.

**Figure 3.**
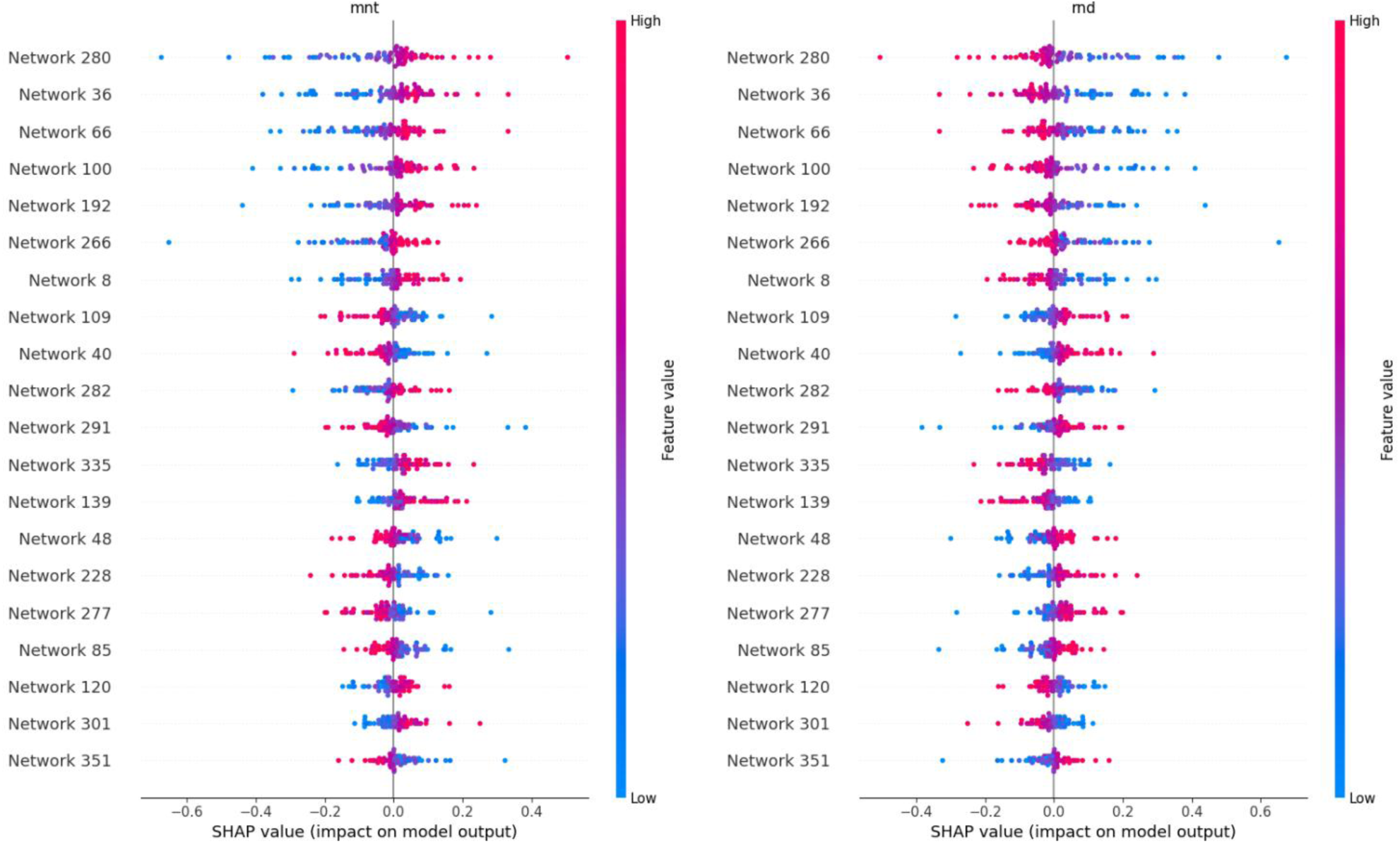
Summary plot containing the distribution of SHAP values of the retrained network for each of the twentieth highest-ranked networks (inputs) for both categories. A higher rank means greater influence on the output computation, either positively (red on the right) or negatively (blue on the right). For example, networks 280 and 36 are the inputs that positively contribute most to the mnt output selection, whereas networks 109 and 40 are the inputs that contribute most to rnd.

All of these brain networks contribute symmetrically across categories: if an input feature contributes positively to the mentalizing (mnt) class, it contributes negatively to the random (rnd) class, and vice versa. This symmetric pattern is expected given the binary structure of the classification task and the additive explanation framework implemented through SHAP.

In this way, brain networks 109, 40, 291, 48, 228, 277, 85, and 351 (sequenced in decreasing order of SHAP values modulus) contribute positively to the rnd class, while the regions 280, 36, 66, 100, 192, 266, 8, 282, 335, 139, 120, and 301 contribute positively to the mnt class. **Table 3** shows the regions that compose these networks, as well as the class they contribute positively to, according to the SHAP analysis.

**Table 3.** Brain networks and respective regions that hold important information for the classification process in this study. Their absence during Shapley values estimation leads to a significant drop in classification performance.

| Network | Region ID | Region name | Lobe | Cortex |
| --- | --- | --- | --- | --- |
| 280 | 280 | Area_OP4-PV_R | Par | Posterior_Opercular |
|  | 296 | Area_PFt_R | Par | Inferior_Parietal |
|  | 328 | Area_PF_Complex_R | Par | Inferior_Parietal |
| 36 | 36 | Area_5m_L | Par | Paracentral_Lobular_and_Mid_Cingulate |
| 66 | 66 | Area_47m_L | Fr | Orbital_and_Polar_Frontal |
| 100 | 100 | Area_OP4-PV_L | Par | Posterior_Opercular |
|  | 116 | Area_PFt_L | Par | Inferior_Parietal |
|  | 148 | Area_PF_Complex_L | Par | Inferior_Parietal |
| 192 | 192 | Area_55b_R | Fr | Premotor |
|  | 275 | Area_Lateral_IntraParietal_dorsal_R | Par | Superior_Parietal |
|  | 311 | Area_TG_dorsal_R | Temp | Lateral_Temporal |
|  | 317 | Area_PHT_R | Temp | Lateral_Temporal |
|  | 324 | Area_IntraParietal_2_R | Par | Inferior_Parietal |
|  | 325 | Area_IntraParietal_1_R | Par | Inferior_Parietal |
|  | 329 | Area_PFm_Complex_R | Par | Inferior_Parietal |
| 266 | 266 | Area_9-46d_R | Fr | Dorsolateral_Prefrontal |
| 8 | 8 | Primary_Motor_Cortex_L | Fr | Somatosensory_and_Motor |
|  | 82 | Area_IFSa_L | Fr | Inferior_Frontal |
|  | 116 | Area_PFt_L | Par | Inferior_Parietal |
|  | 148 | Area_PF_Complex_L | Par | Inferior_Parietal |
|  | 149 | Area_PFm_Complex_L | Par | Inferior_Parietal |
| 109 | 26 | Superior_Frontal_Language_Area_L | Fr | Dorsolateral_Prefrontal |
|  | 44 | Area_6m_anterior_L | Fr | Paracentral_Lobular_and_Mid_Cingulate |
|  | 70 | Area_8B_Lateral_L | Fr | Dorsolateral_Prefrontal |
|  | 104 | RetroInsular_Cortex_L | Par | Early_Auditory |
|  | 109 | Middle_Insular_Area_L | Ins | Insular_and_Frontal_Opercular |
|  | 125 | Auditory_5_Complex_L | Temp | Auditory_Association |
|  | 175 | Auditory_4_Complex_L | Temp | Auditory_Association |
| 40 | 40 | Dorsal_Area_24d_L | Fr | Paracentral_Lobular_and_Mid_Cingulate |
| 282 | 282 | Area_OP2-3-VS_R | Par | Posterior_Opercular |
| 291 | 291 | Anterior_Ventral_Insular_Area_R | Ins | Insular_and_Frontal_Opercular |
| 335 | 302 | Perirhinal_Ectorhinal_Cortex_R | Temp | Medial_Temporal |
|  | 335 | ParaHippocampal_Area_2_R | Temp | Medial_Temporal |
| 139 | 78 | Rostral_Area_6_L | Fr | Premotor |
|  | 79 | Area_IFJa_L | Fr | Inferior_Frontal |
|  | 80 | Area_IFJp_L | Fr | Inferior_Frontal |
|  | 139 | Area_TemporoParietoOccipital_Junction_1_L | Temp | Temporo-Parieto_Occipital_Junction |
| 48 | 48 | Area_Lateral_IntraParietal_ventral_L | Par | Superior_Parietal |
|  | 124 | ParaBelt_Complex_L | Temp | Early_Auditory |
|  | 175 | Auditory_4_Complex_L | Temp | Auditory_Association |
| 228 | 228 | Area_Lateral_IntraParietal_ventral_R | Par | Superior_Parietal |
|  | 304 | ParaBelt_Complex_R | Temp | Early_Auditory |
|  | 355 | Auditory_4_Complex_R | Temp | Auditory_Association |
| 277 | 277 | Inferior_6-8_Transitional_Area_R | Fr | Dorsolateral_Prefrontal |
| 85 | 85 | Area_anterior_9-46v_L | Fr | Dorsolateral_Prefrontal |
| 120 | 120 | Hippocampus_L | Temp | Medial_Temporal |
| 301 | 301 | ProStriate_Area_R | Par | Posterior_Cingulate |
|  | 302 | Perirhinal_Ectorhinal_Cortex_R | Temp | Medial_Temporal |
| 351 | 351 | Area_posterior_47r_R | Fr | Inferior_Frontal |

**Figure 4** shows the spatial expression of these networks. The networks relevant to the rnd class are in shades of blue, while those relevant to the mnt class are in shades of red (brighter colors indicate higher SHAP importance).

**Figure 4.**
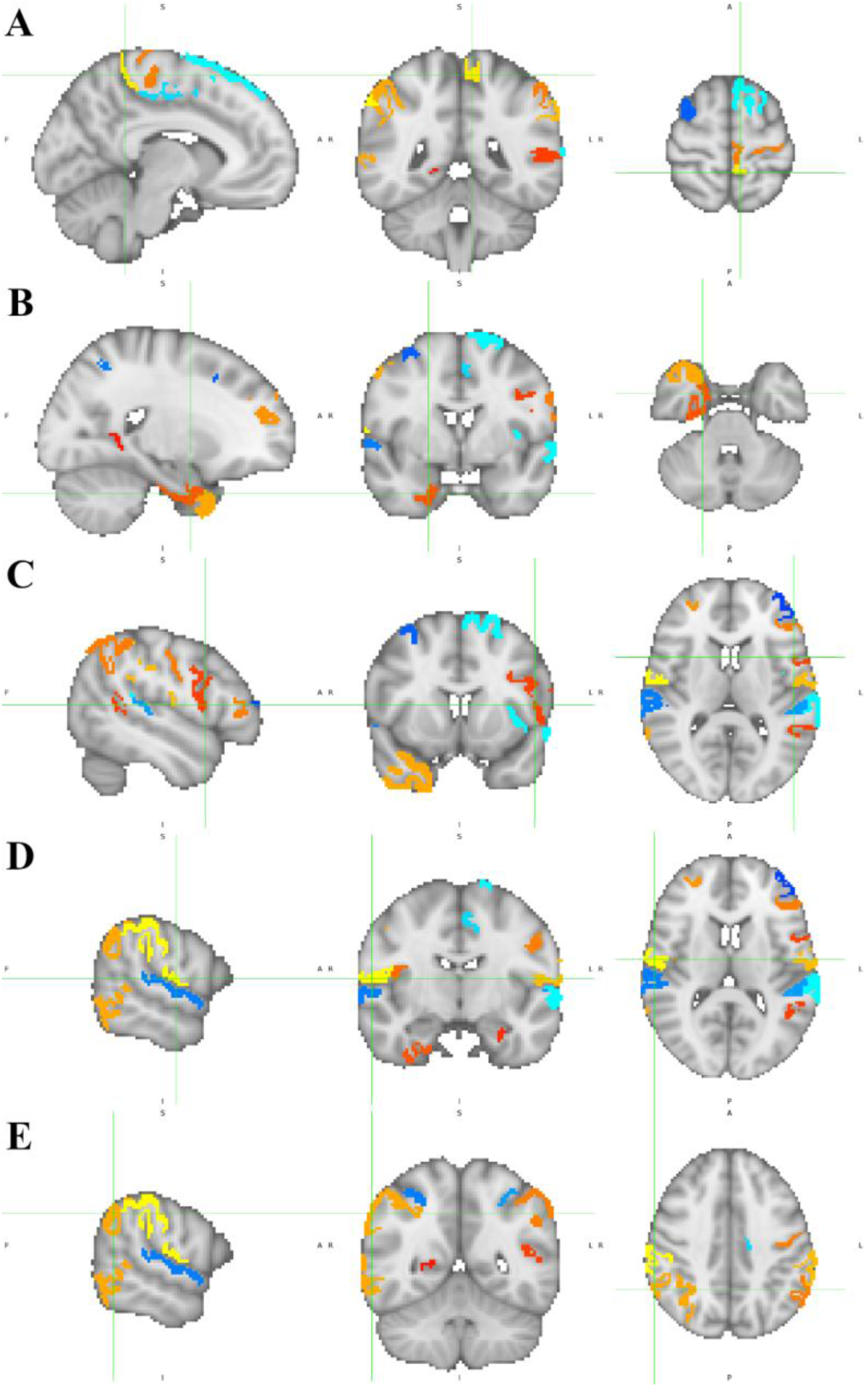
Networks with the twenty highest SHAP values modulus (identified in **Table 3**) depicted over the ICBM152 brain template. Networks that contribute positively to mentalizing (mnt) are in the red-yellow range, networks that contribute positively to random (rnd) are in the dark-light blue range. Lighter colors (closer to red and light blue, for mnt and rnd, respectively) are associated with higher importance. Each pane consists of sagittal, coronal, and axial views of A x = −8, y = −44, z = 62; B x = 22, y = 0, z = −34; C x = −50, y = 10, z = 10; D x = 60, y = −10, z = 8; E x = 50, y = −52, z = 40. MNI152 standard space. Radiological convention. A: anterior; I: inferior; L: left; P: posterior; R: right; S: superior.

These results raise the question of how these networks compare to typical ToM studies in the literature. **Figure 5** shows the overlap between the networks that positively contribute to the mnt class and expected regions based on neuroscientific literature. Neurosynth is used as it returns a map of the regions present on a term-based meta-analysis for the concept “mentalizing”. The regions present in the literature are in green, the mentalizing networks identified with the SNN + SHAP values method are in red, and their overlap is in yellow.

**Figure 5.**
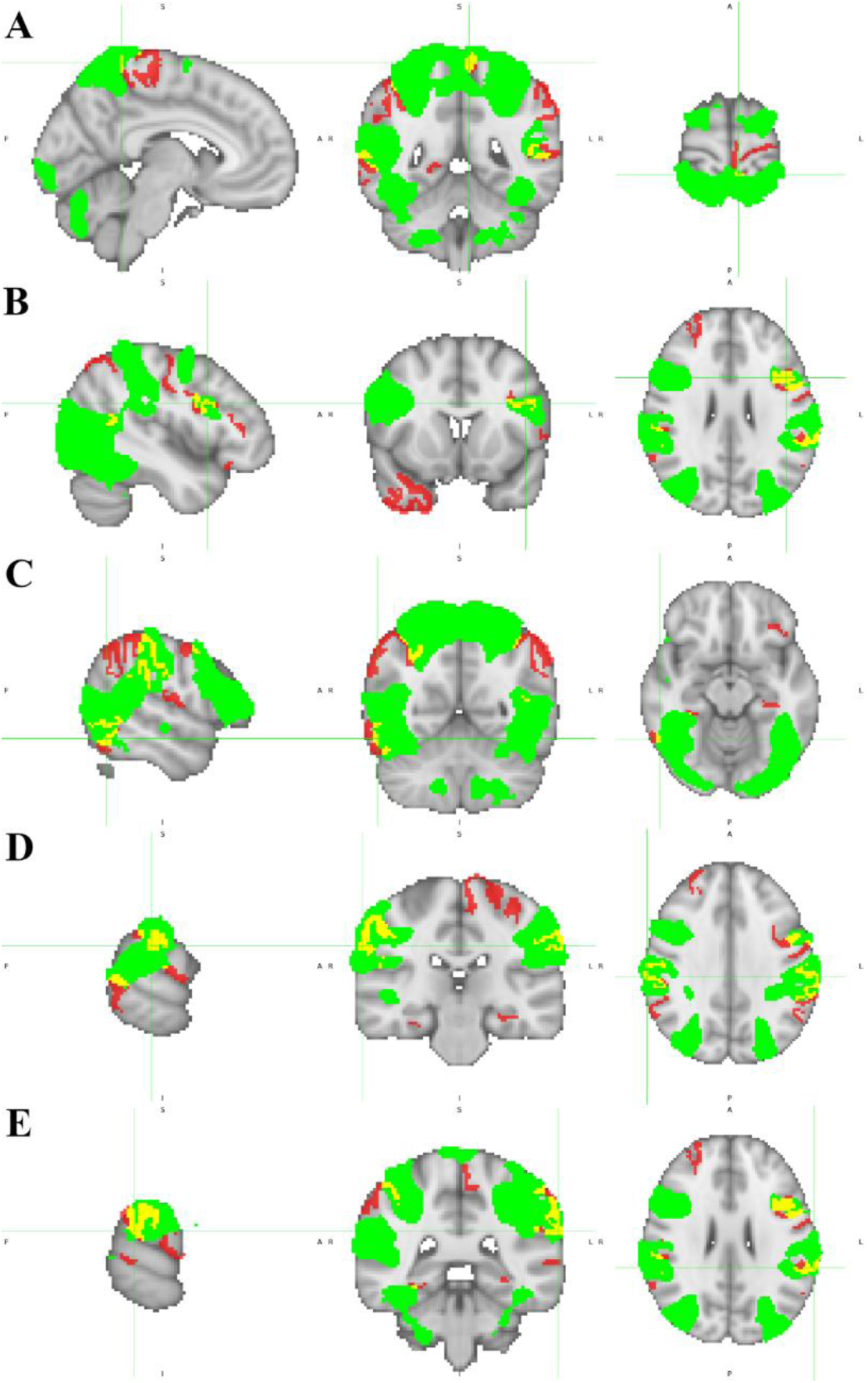
Networks in the top twenty SHAP values that contribute positively to mentalizing overlapped with the GLM-based statistical parametric map obtained from (Dureux et al., 2023). Regions that belong only to the identified networks are in red, areas that are present only in the literature are in green, and regions that are present in both (overlap) are in yellow. Each pane consists of sagittal, coronal, and axial views of A x = −6, y = −46, z = 70; B x = −44, y = 12, z = 26; C x = 56, y = −56, z = −14; D x = 66, y = −26, z = 32; E x = −66, y = −38, z = 26. MNI152 standard space. Radiological convention. A: anterior; I: inferior; L: left; P: posterior; R: right; S: superior.

The regions identified from the GLM in the literature span a total of 47,840 voxels, whereas the networks identified by the proposed method encompass 7,664 voxels. The overlap between the two sets corresponds to 1,910 voxels, resulting in an overlap coefficient of 0.249. Although this value indicates a limited direct spatial overlap, a closer inspection of **Figure 5** reveals that much of this overlap occurs at regions commonly associated with ToM studies. It is also evident that, even with the same threshold as applied in the original paper (*z* > 2.57), the GLM paints massive blobs that include many white-matter voxels. Given the nature of the BOLD signal, it is highly unlikely that activations in white-matter find support in biological activity. Presumably, they are mostly artifacts of spatial smoothing. Furthermore, the atlas chosen for parcellation (MMP1.0) includes only cortical gray-matter. Thus, to make the actual overlap clearer, the GLM map obtained from the literature is masked using the MMP1.0 atlas. All other parameters are kept, which is presented in **Figure 6**.

**Figure 6.**
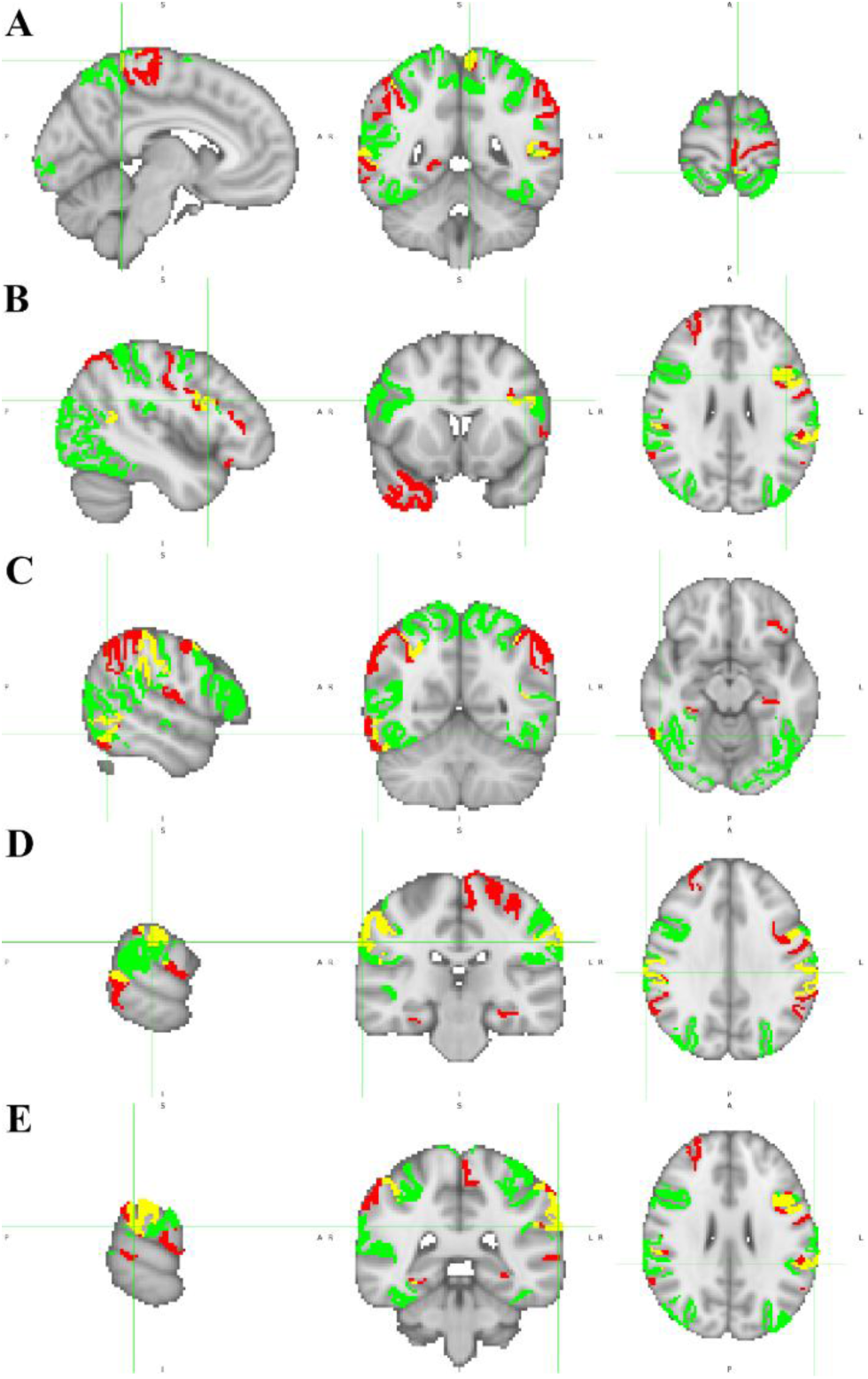
Networks in the top twenty SHAP values that contribute positively to mentalizing overlapped with the GLM-based statistical parametric map obtained from (Dureux et al., 2023) after masking with the MMP1.0 atlas, so that only cortical gray-matter voxels are included for both cases. Regions that belong only to the identified networks are in red, areas that are present only in the literature are in green, and regions that are present in both (overlap) are in yellow. Each pane consists of sagittal, coronal, and axial views of A x = −6, y = −46, z = 70; B x = −44, y = 12, z = 26; C x = 56, y = −56, z = −14; D x = 66, y = −26, z = 32; E x = −66, y = −38, z = 26. MNI152 standard space. Radiological convention. A: anterior; I: inferior; L: left; P: posterior; R: right; S: superior.

This masking reduces the number of voxels in GLM from 47,840 to 17,875. Thus, 29,965 voxels of the GLM-based map are marked as activations with questionable biological support. It is also clearer where the actual overlap between the two methods occurs. The overlap occurs mostly in regions 36 (area 5m of the left paracentral lobular and mid cingulate cortex, postcentral gyrus, pane A of **Figure 6**), 79 (area IFJa of the left inferior frontal cortex, inferior frontal gyrus, pane B of **Figure 6**), 317 (area PHT of the right lateral temporal cortex, inferior temporal gyrus, pane C of **Figure 6**), 328 (area PF complex of the right inferior parietal cortex, supramarginal gyrus, pane D of **Figure 6**), and 148 (area PF complex of the left inferior parietal cortex, supramarginal gyrus, pane E of **Figure 6**). Other relevant overlaps to mention include regions 8, 78, 80, 116, 139, 148, 191, 192, 280, 296, 324, and 329, the complete list of region names and cortices for these is available in Supplementary Table 1.

### 3.4 Identification of Regions within Networks

It is pertinent to identify which regions within the networks the classifier uses to distinguish between mentalizing and random processes that contribute to the response. In this way, Cohen’s D values between regions are computed separately for the mentalizing and random instances. As Cohen’s D measures the distance between the averages, it allows quantification of the relative magnitudes of the signal within each network and, through this, ranks the regions according to their relative contribution to the network’s signal. When paired with the SHAP analysis, this allows the contribution to the network’s signal to be extended to the classifier’s decision. As it is irrelevant to perform this analysis on networks comprising a single region, only the top 5 SHAP-ranked networks comprising at least 2 regions are considered (networks 36 and 66 are unconnected according to the connectivity matrix and thus excluded from this analysis). These results are compiled Table 4 for the mentalizing condition and in Table 5 for the random condition.

**Table 4.**
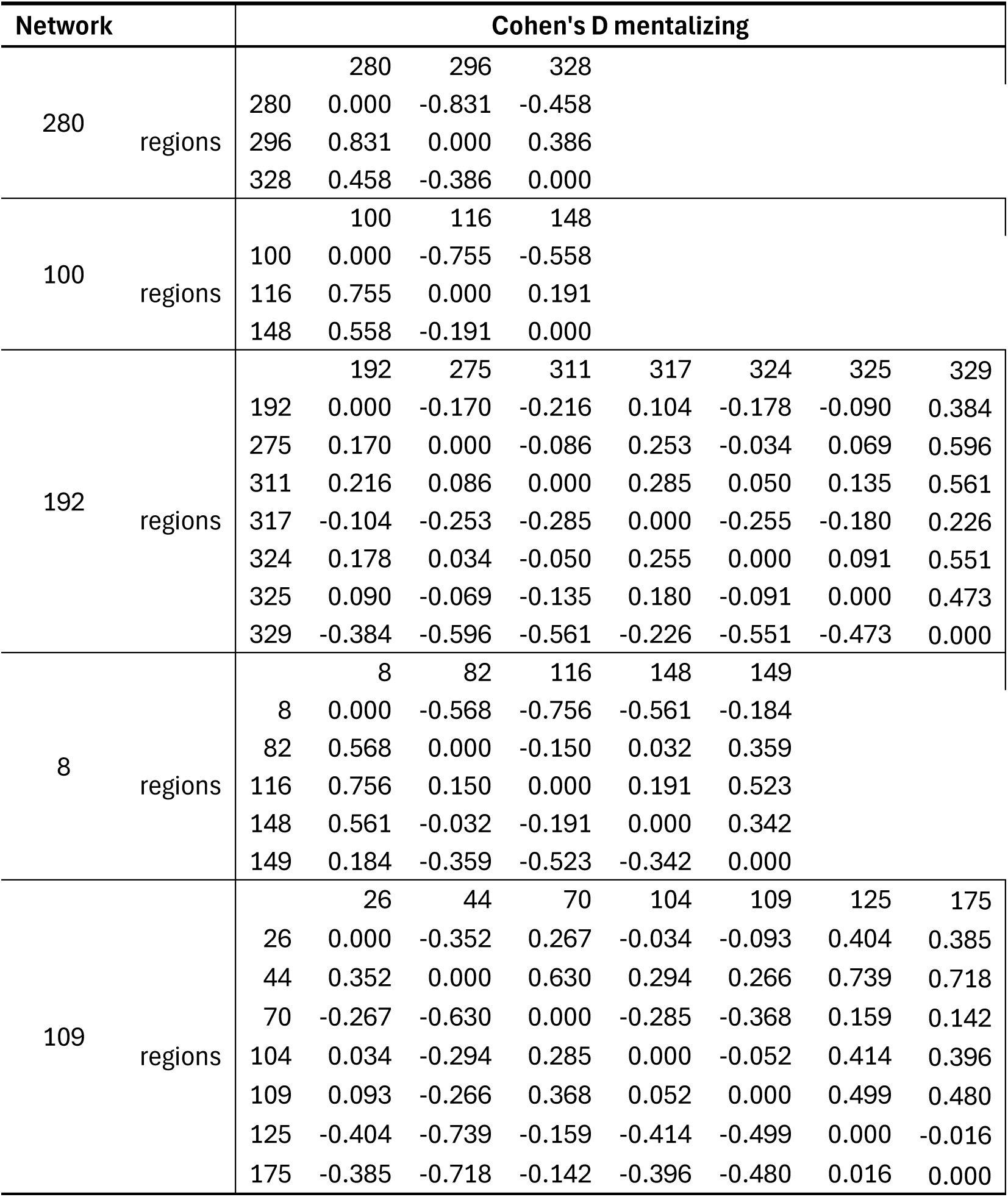
Cohen’s D values for the mentalizing instances across regions within each of the selected networks for this analysis. A lower value indicates the regions have closer signal magnitudes in these instances. A higher value indicates their signals are farther apart.

**Table 5.**
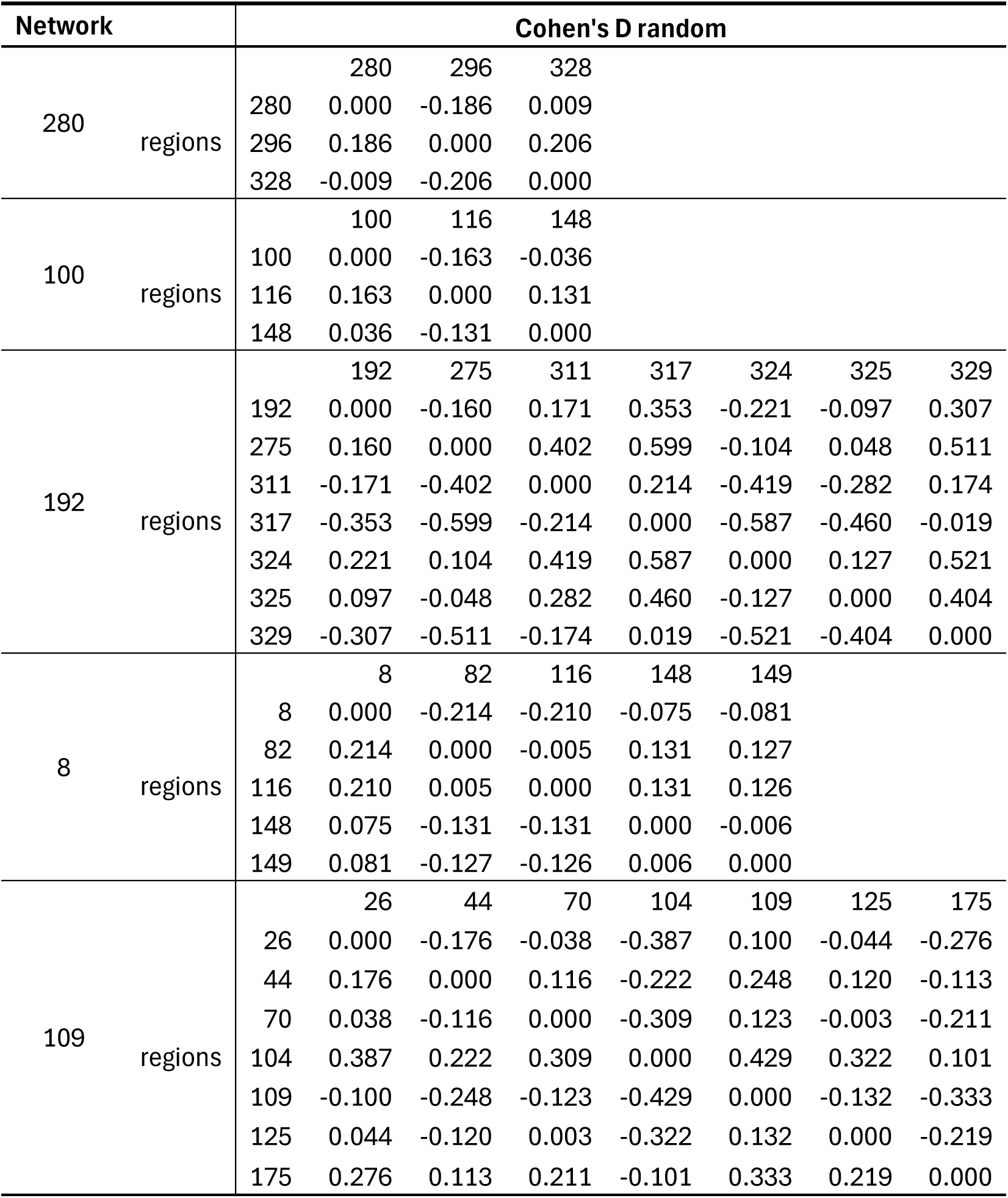
Cohen’s D values for the random instances across regions within each of the selected networks for this analysis. A lower value indicates the regions have closer signal magnitudes in these instances. A higher value indicates their signals are farther apart.

Looking at each network individually across both conditions, in network 280, its regions can be ranked in descending order of signal magnitude for mentalizing as 296, 328, 280, and, interestingly, for the random condition, the order changes to 296, 280, 328, but with these regions being much closer together. For network 100, there is again a clear ranking of its regions (116, 148, 100, in descending order of signal magnitude for mentalizing), and a similar pattern as before is observed for the random condition. For network 192, all regions seem to rank much closer together in the mentalizing condition, except for region 329, which is below the others. This changes slightly for the random condition, with regions 324 and 275 being above the others, and regions 317 and 329 distinguishing themselves as being below the others. For network 8, there is again a clear ranking for mentalizing, with region 116 being above the others, followed by regions 82 and 148, still with a positive average Cohen’s D value, followed by regions 149 and 8; these differences are greatly reduced in the random condition. For network 109, region 44 is clearly the highest for the mentalizing condition, followed by 109 and 104, which are in turn closely followed by region 26, then region 70, with regions 125 and 175 being well below these. For the random condition, these differences are once again greatly reduced. It is important to note that, of these 5 networks, the first 4 (networks 280, 100, 192, and 8) contribute positively to the mentalizing classification, while only network 109 contributes positively towards random. The complete 360-by-360 matrices with the Cohen’s D values for all region pairs are available in Supplementary Tables 3 and 4, for mentalizing and random, respectively.

## 4 Discussion

This study introduces a structurally informed message-passing framework integrated into a shallow neural network classifier. The primary finding is that network-level modeling achieves performance comparable to a region-level baseline while substantially improving biological interpretability. Although the initial global accuracy is slightly lower in the proposed model (87.1% vs. 88.2%), both approaches converge to an identical performance level (84.7%) after pruning and retraining. These results demonstrate that incorporating structural connectivity does not reduce predictive validity, while it shifts the interpretative focus from isolated brain regions toward biologically plausible networks, thereby avoiding the concerns raised by Wen et al. (Wen et al., 2018) (balancing predictive performance with robustness in low-data regimes). In the same way, the proposed framework avoids the risk of capturing spurious or biologically implausible patterns when anatomical constraints are not considered.

Whereas the baseline model identifies individual regions contributing to classification, the proposed approach constrains the propagation of information according to diffusion-based structural connectivity. As a consequence, the explanatory units correspond to structurally grounded networks rather than isolated nodes. This perspective aligns with systems-level approaches in Neuroscience, which conceptualize cognitive processes as emergent properties of distributed interactions across large-scale brain systems. In particular, processes such as Theory of Mind (ToM) are widely understood to rely on coordinated activity across multiple cortical regions rather than on a single localized substrate. The higher post-retraining accuracy observed for the mentalizing condition (95.6%) suggests that structurally organized network recruitment may be especially informative for socially relevant cognitive processes. This is then validated by most of the relevant networks identified by the SHAP analysis, contributing positively to the mentalizing process. In fact, out of the top 20 networks, 12 contribute positively towards mentalizing, including all of the top 7 most relevant networks.

The relatively small sample size used in this study reflects realistic constraints associated with task-based fMRI acquisition, where collecting large cohorts with complex paradigms can be challenging. The fact that the proposed model demonstrates stable convergence and interpretable results under these conditions suggests that the framework is well-suited for typical neuroimaging studies. Nevertheless, evaluating the robustness and generalizability of the approach in larger datasets remains an important direction for future work.

Biological plausibility constitutes a central contribution of the proposed framework. Structurally constrained message-passing restricts the hypothesis space to anatomically feasible configurations, thereby reducing the likelihood that the classifier exploits spurious correlations unrelated to known brain organization. The inclusion of a diagonal term ensures that intra-regional contributions are preserved during signal propagation, while degree normalization mitigates signal dilution effects that could arise when nodes with many connections aggregate information from both informative and non-informative regions. Together, these mechanisms ensure that both local and distributed contributions to the neural signal are meaningfully represented.

Importantly, the spatial distribution of the networks identified as most relevant for classification shows some overlap along with a high number of adjacent regions to those previously implicated in ToM processing. In particular, several of the identified networks involve regions within the temporo-parietal junction (TPJ), superior temporal sulcus (STS), and inferior frontal gyrus (IFG). These regions have repeatedly been reported as core components of the ToM network in prior studies (TPJ: (Lavoie et al., 2016; Ogawa & Matsuyama, 2023; Schurz et al., 2014; Young et al., 2010); STS: (Frith & Frith, 2006; Otsuka et al., 2009; Takahashi et al., 2004); IFG: (Qin et al., 2023)). The correspondence between the networks highlighted by the present framework and regions consistently reported in the literature further supports the robustness and biological relevance of the proposed method. This is more clearly visible in **Figure 6** of the results. The overlap analysis also exposes a key limitation of conventional GLM-based approaches. A substantial proportion of the voxels identified as associated with the mentalizing condition are located in white-matter regions. From a neurophysiological perspective, this is unlikely to reflect task-related activation. The BOLD signal primarily originates from metabolic demand, which is concentrated in gray matter. In contrast, white matter is composed mainly of myelinated axonal fibers, exhibits lower metabolic activity, reduced vascular density, and weaker neurovascular coupling. Consequently, meaningful BOLD signal changes are not expected in these regions.

The presence of apparent white matter activations in GLM results is therefore most plausibly explained by methodological factors, particularly spatial smoothing. While smoothing is commonly applied to improve the signal-to-noise ratio and accommodate inter-subject variability, it also introduces partial-volume effects, effectively blurring signals across tissue boundaries. As a result, strong gray-matter activations can “leak” into adjacent white-matter voxels, leading to biologically implausible findings.

A comparison between the networks identified through the SHAP-based model analysis and the regions previously highlighted in the region-level analysis reported in (Marques dos Santos et al., 2025) provides additional insights. Interestingly, most of the top ten regions identified in the previous region-based analysis do not appear in the most relevant networks in the present study. Only region 125, previously identified as the most important feature for classification, appears within network 109, and region 80, previously ranked fourth, appears within network 139. At the same time, several regions that were previously ranked slightly lower still appear prominently within the identified networks. These include regions 66, 175, 296, 328, 116, 82, 36, 192, 317, and 148, which were ranked 11th, 13th, 17th, 23rd, 32nd, 33rd, 47th, 54th, 58th, and 65th in the region-based SHAP ranking, respectively.

This observation suggests that groups of regions with moderate individual importance may collectively provide stronger information about the distinction between mentalizing and random processes than any single region alone. In other words, network-level organization may capture distributed patterns of activity that are not apparent when features are evaluated independently. Still, this does not fully explain why most of the top-ten regions identified in the previous region-based analysis do not reappear in the most relevant networks. Several mechanisms may explain why certain regions gain prominence in the network-based analysis. One possibility relates to signal dilution: regions that were previously highly informative at the regional level may be structurally connected to many regions that are not directly involved in the task. When signals are aggregated through message-passing, the relevant activity may be partially diluted by baseline signals and noise from these additional regions.

Another possible explanation concerns the role of the hidden layer in the classifier. During training, the hidden nodes may integrate information from multiple structurally defined subnetworks to form higher-level representations that are more informative for classification. In this scenario, the importance of individual regions would depend not only on their own signal but also on how they interact with other regions within the learned representations. A detailed investigation of the signal time series after the message-passing step, as well as an analysis of the hidden-layer activations, would be required to evaluate these hypotheses.

The analysis of Cohen’s D values provides complementary insights into the structure of the network signals. Although these values primarily capture relative differences in signal magnitude and, therefore, cannot definitively determine whether specific regions are completely uninvolved in the task, several patterns nevertheless emerge. In general, the networks exhibit larger Cohen’s D values, i.e., signal magnitudes further apart, for the mentalizing condition than for the random condition.

This observation is particularly notable given that most of the networks identified by the SHAP analysis contribute positively to the mentalizing classification. One possible interpretation is that the larger effect sizes reflect heterogeneous activation within the networks: some regions may be selectively recruited during mentalizing, while others remain closer to baseline. Such variability would lead to larger differences in signal magnitude across regions within the same network. In contrast, during the random condition, signals across the network may remain more uniformly close to baseline levels, indicating high levels of correlation, which translate into lower Cohen’s D values. If confirmed, this pattern would suggest the existence of structurally constrained, yet functionally selective, subnetworks involved in mentalizing. However, this hypothesis requires further validation through analyses of absolute signal magnitudes rather than relying on relative measures alone. Even so, the current analysis already allows ranking regions within each network according to their expected contribution to the model’s classification decisions.

Clinically, the proposed framework may be particularly relevant to the study of neurological and psychiatric disorders, which are increasingly conceptualized as disorders of network connectivity. In conditions such as Alzheimer’s disease and Mild Cognitive Impairment, large-scale connectivity alterations often precede overt structural atrophy. Similarly, disorders including Attention Deficit Hyperactivity Disorder, Autism Spectrum Disorder, Bipolar Disorder, and Schizophrenia are increasingly described as dysconnectivity syndromes (Du et al., 2018). In such contexts, analytical frameworks that explicitly model interactions among structurally connected regions may yield more informative biomarkers than approaches that treat brain regions in isolation. By integrating anatomical constraints with interpretable machine learning methods, the present approach therefore offers a promising direction for studying both normative cognitive processes and network-level pathology.

An additional extension of the present framework concerns identifying regions that are active under both experimental conditions. A key limitation of classifier explanation methods, such as SHAP as employed in this study, is that they highlight features that differentiate between classes. Because the classifier is optimized to minimize prediction error, features that are similarly active in both conditions will tend to receive low importance scores. A similar limitation exists in traditional statistical analyses based on contrasts between conditions. In such approaches, regions engaged in both processes may be overlooked, even if they are functionally relevant to the task. When studying complex cognitive functions such as mentalizing, however, it is often desirable to identify all regions involved in the process rather than only those that differentiate it from a control condition. In the present framework, such regions may be detected by combining SHAP values with measures of signal magnitude. Regions that are highly active but show low SHAP importance may represent task-relevant components common to both conditions, whereas regions with both low SHAP values and low signal magnitude are more likely to be truly irrelevant to the task.

The adjacency matrix is pivotal within the method proposed here. The adjacency matrix must closely represent the biological network, internalizing in its parameters the tracts and fascicles that connect brain regions, faithfully capturing its small-world structure. This study integrates several works that used anatomy-driven deterministic tractography on diffusion-weighted images from 10 subjects (Baker et al., 2018). Human brain variability within normality may not have been fully accounted for, and population-based adjacency matrices may be designed, yielding different results. Structural impairments may also be modeled by the adjacency matrix. All these branches represent avenues for future work and comparison with the present results and interpretation.

In summary, the present study demonstrates that incorporating structurally constrained message-passing into an SNN classifier enables biologically accurate and interpretable brain decoding without compromising classification performance. By constraining information flow according to anatomical connectivity, the proposed framework shifts interpretation from isolated regions toward distributed brain networks that more closely reflect the organization of cognitive processes described in Neuroscience. The results show that network-level modeling achieves performance comparable to region-based approaches while highlighting subnetworks consistent with prior literature on Theory of Mind. Moreover, the combination of structural constraints and explainability methods such as SHAP enables the identification of networks and constituent regions that contribute to classification, providing insights into both differentiating and potentially shared components of task-related brain activity. Beyond the present paradigm, this framework offers a promising avenue for improving the interpretability of machine learning models applied to fMRI data, particularly in the study of cognitive processes and disorders characterized by alterations in large-scale brain connectivity.

## 5 Author Contributions

All authors have read and agreed to the published version of the manuscript.

Bruno Direito: Conceptualization, Funding acquisition, Investigation, Methodology, Supervision, Writing – review & editing. José Paulo Marques dos Santos: Conceptualization, Formal Analysis, Funding acquisition, Investigation, Methodology, Project administration, Software, Validation, Writing – original draft, Writing – review & editing. Luis Paulo Reis: Funding acquisition, Methodology, Software, Supervision, Writing – review & editing. José Diogo Marques dos Santos: Conceptualization, Data curation, Formal Analysis, Investigation, Methodology, Resources, Software, Validation, Visualization, Writing – original draft, Writing – review & editing. Maria Beatriz Ramos: Formal Analysis, Investigation, Resources, Software, Validation, Writing – review & editing.

## Data Availability Statement

Generated Statement: Publicly available datasets were analyzed in this study. This data can be found here: HCP-Young Adult 2025 at https://balsa.wustl.edu/project?project=HCP_YA.

## References

Adolphs, R. (2001). The neurobiology of social cognition. Current Opinion in Neurobiology, 11(2), 231–239. 10.1016/S0959-4388(00)00202-6

Baker, C. M., Burks, J. D., Briggs, R. G., Conner, A. K., Glenn, C. A., Sali, G., McCoy, T. M., Battiste, J. D., O’Donoghue, D. L., & Sughrue, M. E. (2018). A Connectomic Atlas of the Human Cerebrum-Chapter 1: Introduction, Methods, and Significance. Operative Neurosurgery, 15(suppl_1), S1–S9. 10.1093/ons/opy253

Baliga, V. B., Armstrong, M. S., Press, E. R., Bonnet-Lebrun, A.-S., & Sciaini, M. (2023). pathviewr: Wrangle, Analyze, and Visualize Animal Movement Data (v. 1.1.7). In https://cran.r-project.org/package=pathviewr

Bassett, D. S., & Sporns, O. (2017). Network neuroscience. Nature Neuroscience, 20(3), 353–364. 10.1038/nn.4502

Briggs, R. G., Conner, A. K., Baker, C. M., Burks, J. D., Glenn, C. A., Sali, G., Battiste, J. D., O’Donoghue, D. L., & Sughrue, M. E. (2018). A Connectomic Atlas of the Human Cerebrum-Chapter 18: The Connectional Anatomy of Human Brain Networks. Operative Neurosurgery, 15(suppl_1), S470–S480. 10.1093/ons/opy272

Briggs, R. G., Conner, A. K., Rahimi, M., Sali, G., Baker, C. M., Burks, J. D., Glenn, C. A., Battiste, J. D., & Sughrue, M. E. (2018). A Connectomic Atlas of the Human Cerebrum-Chapter 14: Tractographic Description of the Frontal Aslant Tract. Operative Neurosurgery, 15(suppl_1), S444–S449. 10.1093/ons/opy268

Briggs, R. G., Conner, A. K., Sali, G., Rahimi, M., Baker, C. M., Burks, J. D., Glenn, C. A., Battiste, J. D., & Sughrue, M. E. (2018a). A Connectomic Atlas of the Human Cerebrum-Chapter 16: Tractographic Description of the Vertical Occipital Fasciculus. Operative Neurosurgery, 15(suppl_1), S456–S461. 10.1093/ons/opy270

Briggs, R. G., Conner, A. K., Sali, G., Rahimi, M., Baker, C. M., Burks, J. D., Glenn, C. A., Battiste, J. D., & Sughrue, M. E. (2018b). A Connectomic Atlas of the Human Cerebrum-Chapter 17: Tractographic Description of the Cingulum. Operative Neurosurgery, 15(suppl_1), S462–S469. 10.1093/ons/opy271

Briggs, R. G., Rahimi, M., Conner, A. K., Sali, G., Baker, C. M., Burks, J. D., Glenn, C. A., Battiste, J. D., & Sughrue, M. E. (2018). A Connectomic Atlas of the Human Cerebrum-Chapter 15: Tractographic Description of the Uncinate Fasciculus. Operative Neurosurgery, 15(suppl_1), S450–S455. 10.1093/ons/opy269

Call, J., & Tomasello, M. (2008). Does the chimpanzee have a theory of mind? 30 years later. Trends in Cognitive Sciences, 12(5), 187–192. 10.1016/j.tics.2008.02.010

Chauhan, A. S., Singh, R., Priyadarshi, N., Twala, B., Suthar, S., & Swami, S. (2024). Unleashing the power of advanced technologies for revolutionary medical imaging: Pioneering the healthcare frontier with artificial intelligence. Discover Artificial Intelligence, 4(1), 58. 10.1007/s44163-024-00161-0

Conner, A. K., Briggs, R. G., Rahimi, M., Sali, G., Baker, C. M., Burks, J. D., Glenn, C. A., Battiste, J. D., & Sughrue, M. E. (2018a). A Connectomic Atlas of the Human Cerebrum-Chapter 10: Tractographic Description of the Superior Longitudinal Fasciculus. Operative Neurosurgery, 15(suppl_1), S407–S422. 10.1093/ons/opy264

Conner, A. K., Briggs, R. G., Rahimi, M., Sali, G., Baker, C. M., Burks, J. D., Glenn, C. A., Battiste, J. D., & Sughrue, M. E. (2018b). A Connectomic Atlas of the Human Cerebrum-Chapter 12: Tractographic Description of the Middle Longitudinal Fasciculus. Operative Neurosurgery, 15(suppl_1), S429–S435. 10.1093/ons/opy266

Conner, A. K., Briggs, R. G., Sali, G., Rahimi, M., Baker, C. M., Burks, J. D., Glenn, C. A., Battiste, J. D., & Sughrue, M. E. (2018). A Connectomic Atlas of the Human Cerebrum-Chapter 13: Tractographic Description of the Inferior Fronto-Occipital Fasciculus. Operative Neurosurgery, 15(suppl_1), S436–S443. 10.1093/ons/opy267

Decety, J., & Keenan, J. P. (2006). Social Neuroscience: A new journal. Social Neuroscience, 1(1), 1–4. 10.1080/17470910600683549

Doran, D., Schulz, S., & Besold, T. R. (2018). What Does Explainable AI Really Mean? A New Conceptualization of Perspectives. In T. R. Besold & O. Kutz (Eds.), Proceedings of the 1st International Workshop on Comprehensibility and Explanation in AI and ML, CEX 2017 (Vol. 2071). CEUR Workshop Proceedings.

Du, Y., Fu, Z., & Calhoun, V. D. (2018). Classification and prediction of brain disorders using functional connectivity: Promising but challenging [Review]. Frontiers in Neuroscience, 12. 10.3389/fnins.2018.00525

Dureux, A., Zanini, A., Selvanayagam, J., Menon, R. S., & Everling, S. (2023). Gaze patterns and brain activations in humans and marmosets in the Frith-Happé theory-of-mind animation task. Elife, 12, e86327. 10.7554/eLife.86327

Elam, J. S., Glasser, M. F., Harms, M. P., Sotiropoulos, S. N., Andersson, J. L. R., Burgess, G. C., Curtiss, S. W., Oostenveld, R., Larson-Prior, L. J., Schoffelen, J.-M., Hodge, M. R., Cler, E. A., Marcus, D. M., Barch, D. M., Yacoub, E., Smith, S. M., Ugurbil, K., & Van Essen, D. C. (2021). The Human Connectome Project: A retrospective. Neuroimage, 244, 118543. 10.1016/j.neuroimage.2021.118543

Farahani, F. V., Fiok, K., Lahijanian, B., Karwowski, W., & Douglas, P. K. (2022). Explainable AI: A review of applications to neuroimaging data [Systematic Review]. Frontiers in Neuroscience, Volume 16 - 2022. 10.3389/fnins.2022.906290

Frith, C. D., & Frith, U. (2006). The neural basis of mentalizing. Neuron, 50(4), 531–534. 10.1016/j.neuron.2006.05.001

Glasser, M. F., Coalson, T. S., Robinson, E. C., Hacker, C. D., Harwell, J., Yacoub, E., Ugurbil, K., Andersson, J., Beckmann, C. F., Jenkinson, M., Smith, S. M., & Van Essen, D. C. (2016). A multi-modal parcellation of human cerebral cortex. Nature, 536(7615), 171–178. 10.1038/nature18933

Haykin, S. (2009). Neural Networks and Learning Machines (3rd ed.). Prentice Hall.

Heider, F., & Simmel, M. (1944). An experimental study of apparent behavior. The American Journal of Psychology, 57(2), 243–259. 10.2307/1416950

Lavoie, M.-A., Vistoli, D., Sutliff, S., Jackson, P. L., & Achim, A. M. (2016). Social representations and contextual adjustments as two distinct components of the Theory of Mind brain network: Evidence from the REMICS task. Cortex, 81, 176–191. 10.1016/j.cortex.2016.04.017

Limas, M. C., Meré, J. B. O., Marcos, A. G., Ascacibar, F. J. M. d. P., Espinoza, A. V. P., Elías, F. A., & Ramos, J. M. P. (2014). AMORE: A MORE flexible neural network package (0.2-15). In (Version 0.2-15) http://cran.nexr.com/web/packages/AMORE/index.html

Lundberg, S. M., & Lee, S.-I. (2017). A unified approach to interpreting model predictions Advances in Neural Information Processing Systems 30 (NIPS 2017), Long Beach (CA), USA. https://proceedings.neurips.cc/paper/2017/file/8a20a8621978632d76c43dfd28b67767-Paper.pdf

Marques dos Santos, J. D., & Marques dos Santos, J. P. (2022). Towards XAI: Interpretable Shallow Neural Network Used to Model HCP’s fMRI Motor Paradigm Data. In I. Rojas, O. Valenzuela, F. Rojas, L. J. Herrera, & F. Ortuño (Eds.), Bioinformatics and Biomedical Engineering. Lecture Notes in Computer Science (Vol. 13347, pp. 260–274). Springer International Publishing. 10.1007/978-3-031-07802-6_22

Marques dos Santos, J. D., & Marques dos Santos, J. P. (2023). Path weights analyses in a shallow neural network to reach Explainable Artificial Intelligence (XAI) of fMRI data. In G. Nicosia, V. Ojha, E. La Malfa, G. La Malfa, P. Pardalos, G. Di Fatta, G. Giuffrida, & R. Umeton (Eds.), Machine Learning, Optimization, and Data Science. Lecture Notes in Computer Science (Vol. 13811, pp. 417–431). Springer International Publishing. 10.1007/978-3-031-25891-6_31

Marques dos Santos, J. D., Reis, L. P., & Marques dos Santos, J. P. (2025). Decoding mental states in social cognition: Insights from explainable artificial intelligence on HCP fMRI data. Machine Learning and Knowledge Extraction, 7(1), 17. 10.3390/make7010017

Marques dos Santos, J. P., & Marques dos Santos, J. D. (2024). XAI (Explainable Artificial Intelligence) in Neuromarketing / Consumer Neuroscience: An fMRI study on brand perception. Frontiers in Human Neuroscience, 18. 10.3389/fnhum.2024.1305164

Mohammadi, H., & Karwowski, W. (2025). Graph Neural Networks in Brain Connectivity Studies: Methods, Challenges, and Future Directions. Brain Sciences, 15(1), 17.

Norman, G. J., Cacioppo, J. T., & Berntson, G. G. (2010). Social neuroscience. Wiley Interdisciplinary Reviews: Cognitive Science, 1(1), 60–68. 10.1002/wcs.29

Ogawa, K., & Matsuyama, Y. (2023). Heterogeneity of social cognition between visual perspective-taking and theory of mind in the temporo-parietal junction. Neuroscience Letters, 807, 137267. 10.1016/j.neulet.2023.137267

Otsuka, Y., Osaka, N., Ikeda, T., & Osaka, M. (2009). Individual differences in the theory of mind and superior temporal sulcus. Neuroscience Letters, 463(2), 150–153. 10.1016/j.neulet.2009.07.064

Pereira, F., Mitchell, T. M., & Botvinick, M. (2009). Machine learning classifiers and fMRI: a tutorial overview. Neuroimage, 45(1 Suppl), S199–S209. 10.1016/j.neuroimage.2008.11.007

Premack, D., & Woodruff, G. (1978). Does the chimpanzee have a theory of mind? Behavioral and Brain Sciences, 1(4), 515–526. 10.1017/S0140525X00076512

Qin, X., Huang, H., Liu, Y., Zheng, F., Zhou, Y., & Wang, H. (2023). Increased functional connectivity involving the parahippocampal gyrus in patients with schizophrenia during Theory of Mind processing: A psychophysiological interaction study. Brain Sciences, 13(4), 692. 10.3390/brainsci13040692

Repple, J., Gruber, M., Mauritz, M., de Lange, S. C., Winter, N. R., Opel, N., Goltermann, J., Meinert, S., Grotegerd, D., Leehr, E. J., Enneking, V., Borgers, T., Klug, M., Lemke, H., Waltemate, L., Thiel, K., Winter, A., Breuer, F., Grumbach, P.,…Dannlowski, U. (2023). Shared and Specific Patterns of Structural Brain Connectivity Across Affective and Psychotic Disorders. Biological Psychiatry, 93(2), 178–186. 10.1016/j.biopsych.2022.05.031

Roscher, R., Bohn, B., Duarte, M. F., & Garcke, J. (2020). Explainable machine learning for scientific insights and discoveries. IEEE Access, 8, 42200–42216. 10.1109/ACCESS.2020.2976199

Sali, G., Briggs, R. G., Conner, A. K., Rahimi, M., Baker, C. M., Burks, J. D., Glenn, C. A., Battiste, J. D., & Sughrue, M. E. (2018). A Connectomic Atlas of the Human Cerebrum-Chapter 11: Tractographic Description of the Inferior Longitudinal Fasciculus. Operative Neurosurgery, 15(suppl_1), S423–S428. 10.1093/ons/opy265

Saxe, R. (2006). Four Brain Regions for One Theory of Mind? In J. T. Cacioppo, P. S. Visser, & C. L. Pickett (Eds.), Social Neuroscience: People Thinking About Thinking People (pp. 83–101). MIT Press.

Schurz, M., Radua, J., Aichhorn, M., Richlan, F., & Perner, J. (2014). Fractionating theory of mind: A meta-analysis of functional brain imaging studies. Neuroscience & Biobehavioral Reviews, 42, 9–34. 10.1016/j.neubiorev.2014.01.009

Shamay-Tsoory, S. G., Aharon-Peretz, J., & Perry, D. (2009). Two systems for empathy: a double dissociation between emotional and cognitive empathy in inferior frontal gyrus versus ventromedial prefrontal lesions. Brain, 132(3), 617–627. 10.1093/brain/awn279

Shrikumar, A., Greenside, P., & Kundaje, A. (2017, 06–11 Aug, 2017). Learning important features through propagating activation differences. 34th International Conference on Machine Learning, PMLR.

Sievers, B., & Thornton, M. A. (2024). Deep social neuroscience: The promise and peril of using artificial neural networks to study the social brain. Social Cognitive and Affective Neuroscience, 19(1). 10.1093/scan/nsae014

Singer, T., & Tusche, A. (2014). Understanding Others: Brain Mechanisms of Theory of Mind and Empathy. In P. W. Glimcher & E. Fehr (Eds.), Neuroeconomics: Decision Making and the Brain (2nd ed., pp. 513–532). Academic Press. 10.1016/B978-0-12-416008-8.00027-9

Takahashi, H., Yahata, N., Koeda, M., Matsuda, T., Asai, K., & Okubo, Y. (2004). Brain activation associated with evaluative processes of guilt and embarrassment: an fMRI study. Neuroimage, 23(3), 967–974. 10.1016/j.neuroimage.2004.07.054

Van Essen, D. C., & Glasser, M. F. (2016). The Human Connectome Project: Progress and Prospects. Cerebrum: the Dana Forum on Brain Science, 2016, cer–10–16.

Van Essen, D. C., Smith, S. M., Barch, D. M., Behrens, T. E. J., Yacoub, E., & Ugurbil, K. (2013). The WU-Minn Human Connectome Project: An overview. Neuroimage, 80, 62–79. 10.1016/j.neuroimage.2013.05.041

van Mourik, F., Jutte, A., Berendse, S. E., Bukhsh, F. A., & Ahmed, F. (2024). Tertiary review on explainable artificial intelligence: Where do we stand? Machine Learning and Knowledge Extraction, 6(3), 1997–2017. 10.3390/make6030098

Vilone, G., & Longo, L. (2021). Classification of explainable artificial intelligence methods through their output formats. Machine Learning and Knowledge Extraction, 3(3), 615–661. 10.3390/make3030032

Wen, D., Wei, Z., Zhou, Y., Li, G., Zhang, X., & Han, W. (2018). Deep learning methods to process fMRI data and their application in the diagnosis of cognitive impairment: A brief overview and our opinion [Opinion]. Frontiers in Neuroinformatics, 12. 10.3389/fninf.2018.00023

Young, L., Dodell-Feder, D., & Saxe, R. (2010). What gets the attention of the temporo-parietal junction? An fMRI investigation of attention and theory of mind. Neuropsychologia, 48(9), 2658–2664. 10.1016/j.neuropsychologia.2010.05.012

