## supplementary materials for "Structurally Restricted Message-Passing within Shallow Architectures for Explainable Network-Level Brain Decoding on Small Cohorts"

### *Supplementary Material*

#### 1 HCP MMP1.0 Regions

Supplementary Table 1 is the complete list of all 360 regions used in the study, along with their ID, region abbreviation, region name, hemisphere, lobe, cortex ID, cortex, x, y, and z coordinates, and volume.

N.B. Because MS Excel files are not compatible with bioRxiv, this table will be available only after final publication.

#### 2 Connectivity Matrix

Supplementary Table 2 is the connectivity matrix obtained from the literature after adding the diagonal term to account for intra-regional contributions during the message-passing.

N.B. Because MS Excel files are not compatible with bioRxiv, this table will be available only after final publication.

#### 3 SHAP

Supplementary Figure 1 depicts the full lists of the Shapley values for mentalizing and random conditions in the retrained network. The networks (features) are ranked by their importance. The more important networks for the model's final prediction are plotted closer to the top.

#### 4 Cohen's D values

Supplementary Table 3 is the complete matrix of Cohen's D values for all possible region pairs for the mentalizing condition. Supplementary Table 4 is the complete matrix of Cohen's D values for all possible region pairs for the random condition.

N.B. Because MS Excel files are not compatible with bioRxiv, this table will be available only after final publication.

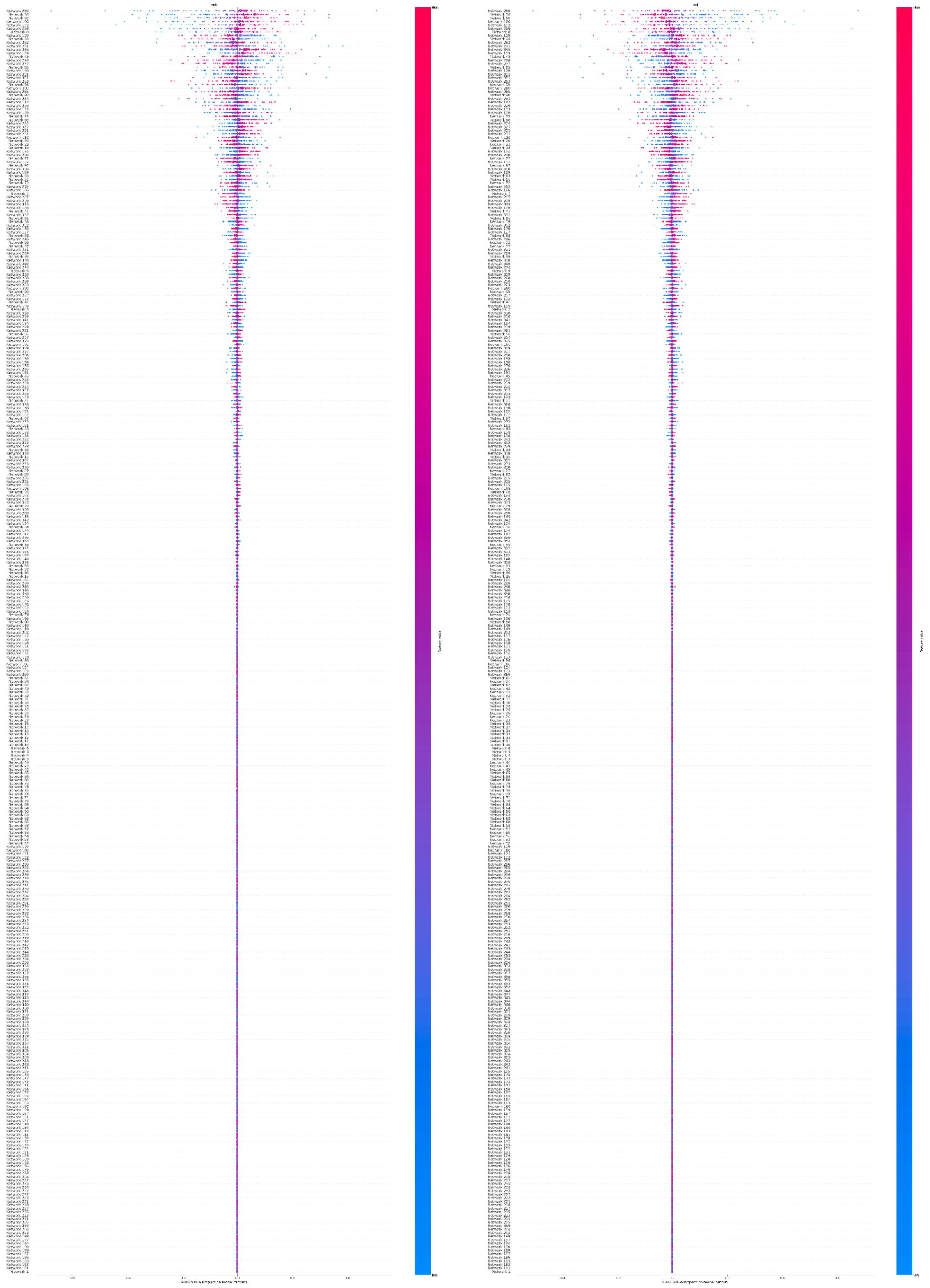

**Supplementary Figure 1** Plot containing the distribution of SHAP values of the retrained network for all networks (inputs) for both category. Higher rank means more influence in the output computation either positively (red on the right), or negatively (blue on the right).
